# A mobile optical coherence microscope for studying aquatic organisms in- and outside the laboratory

**DOI:** 10.64898/2026.07.27.740915

**Authors:** Ling Wang, Samuel Davis, Tabea Quilitz, Thomas Beavis, Michael Bonadonna, Mantha Lamprousi, Roberto Montanari, Fabian Ruperti, Anniek Stokkermans, Tina Wiegand, Victoria Alicia Witte, Alejandro Gil Ortiz, Michael Dorrity, Jan Siemens, Jacob Musser, Detlev Arendt, Nikolaus Leisch, Flora Vincent, Aissam Ikmi, Robert Prevedel

**Affiliations:** Cell Biology and Biophysics Unit, European Molecular Biology Laboratory, Heidelberg, Germany; Developmental Biology Unit, European Molecular Biology Laboratory, Heidelberg, Germany; Mobile Laboratory Facilities, European Molecular Biology Laboratory, Heidelberg, Germany; Molecular Systems Biology Unit, European Molecular Biology Laboratory, Heidelberg, Germany; Institute of Pharmacology, Heidelberg University, Heidelberg, Germany; Molecular Medicine Partnership Unit (MMPU), European Molecular Biology Laboratory; Electronics Workshop, European Molecular Biology Laboratory, Heidelberg, Germany; Department of Molecular, Cellular and Developmental Biology, Yale University; New Haven, 6511, USA

**Author notes:** Correspondence and Lead Contact: R.P. College of Optical and Electronic Technology, China Jiliang University, Hangzhou, Zhejiang, China (Ling Wang).

## Abstract

High-resolution, three-dimensional imaging of live organisms has largely depended on laboratory-bound microscopes, limiting quantitative analysis of morphology and dynamics to species that survive transport and thrive under lab-controlled conditions. Here we present a mobile optical coherence microscopy (OCM) platform that overcomes this constraint, delivering ∼2.5 µm axial resolution, label-free, volumetric imaging of live aquatic organisms in both laboratory and remote field environments. We demonstrate the platform across a broad range of aquatic organisms from the lab and field, spanning multiple phyla - including cnidarians, poriferans, annelids, arthropods and echinoderms – resolving internal anatomy, tissue boundaries and organismal morphology at micrometer scale without fixation, fluorescent labeling, or specialized sample preparation. High-speed acquisition, with up to 250 kHz A-line rate and 7.7 Hz volume rate, further enabled morphodynamic imaging of live biological processes, including cellular aggregate motility, embryonic cell division, and organ-level peristaltic dynamics in intact, living animals. To demonstrate field deployability, the platform was operated during the EMBL TREC pan-European expedition, enabling on-site, label-free imaging of marine organisms and plankton immediately upon collection. By decoupling high-resolution volumetric imaging from fixed laboratory settings, mobile OCM opens a path toward field-ready quantitative morphological and dynamic phenotyping of aquatic life.

## Introduction

Aquatic environments, from oceans to freshwater ecosystems, harbor much of life’s phylogenetic and anatomical diversity, yet experimental analysis of organismal morphology, physiology, and development remains largely confined to species that can be transported, maintained and imaged under laboratory conditions^1,2^. For many aquatic organisms, from marine plankton to soft-bodied invertebrates, the biological state of interest can be altered or lost during collection, preservation, transport or laboratory habituation, before conventional imaging can begin^3–5^. Meanwhile, the growing urgency of climate change, biodiversity loss, and ocean monitoring demands tools that can characterize organismal biology at the sampling site, in real time, across a broader range of aquatic life^6,7^. Existing high-resolution, three-dimensional imaging methods have so far been unable to meet this demand: existing modalities either lack the spatio-temporal resolution to resolve tissue-level structure and function, require fluorescent labeling that can be impractical or time-consuming for non-model organisms, or depend on laboratory infrastructure such as vibration isolation and precise temperature control incompatible with field deployment.

Clinical imaging approaches like ultrasound, CT, and MRI, lack the resolution to resolve tissue microstructure in small organisms, and MRI is incompatible with live aquatic specimens^8–11^. Confocal^12^ and light-sheet^13^ microscopy offer the necessary resolution but require fluorescent labeling, complex sample mounting, and laboratory infrastructure, often precluding their straightforward use for most non-model species and for field deployment, although notable exceptions do exist^5,14^. In contrast, optical coherence tomography (OCT) provides label-free, near-infrared contrast with high acquisition speed and millimeter-scale penetration depth, and has been applied to aquatic invertebrates including scallop, coral, and jellyfish^15–18^. However, prior implementations used low-magnification optics and stationary, laboratory-bound systems, limiting both lateral resolution and accessibility to field environments. So far, an imaging modality that combines micrometer-scale three-dimensional live imaging with label-free contrast and high acquisition speed, and within a form factor compatible with ready deployment beyond the laboratory has not been realized.

Optical coherence microscopy (OCM), an advanced adaptation of optical coherence tomography (OCT), integrates high magnification image forming optics, and thus can offer superior resolution that can enable more detailed visualization of cellular and subcellular structures^19–21^. In this work, we present a mobile OCM platform optimized for high-resolution, label-free imaging and biometry of live aquatic organisms spanning multiple phyla - including cnidarians^22^ (*Nematostella vectensis, Hydra*), poriferans^22^ (*Spongilla lacustris*), annelids (*Platynereis dumerilii*), and arthropods - as well as marine plankton and a vertebrate model organism (*Danio rerio*), all imaged under near-native conditions. Importantly, our OCM system was fully miniaturized and tailored for operation outside conventional laboratory environments. The robustness and versatility of the mobile OCM platform were demonstrated during multiple field deployments, including the EMBL TREC expedition^23^. In these settings, the system enabled non-invasive, high-resolution imaging of a wide range of marine and freshwater species under near-native conditions, illustrating its suitability for in situ biological imaging and quantitative biometry in diverse field environments.

## Results

### Mobile OCM system design and performance

To meet the requirements of high resolution and rapid imaging of live aquatic organisms, we developed a spectral-domain OCM system optimized for high-resolution, label-free imaging in both laboratory and field settings. Fig. 1a shows a schematic diagram of the OCM system which is built around a fiber coupler-based Michelson interferometer illuminated by a superluminescent diode (SLD) with a ∼160 nm bandwidth centered near 850 nm, yielding a measured axial resolution of ∼2.5 µm in water. The sample arm employs a galvo mirror pair conjugated to the back focal plane of interchangeable objectives (2×, 4×, and 10×, NA 0.06– 0.30) via a 4f scanning configuration, providing lateral resolutions between 1.4 and 7.0 µm, respectively, depending on the objective lens (see Methods). A custom spectrometer equipped with a high-speed line-scan camera (Octoplus, Teledyne e2V) enables acquisition at up to 250 kHz A-line rates, supporting volumetric imaging at up to 7.7 Hz over 180 × 180 lateral pixels. At this speed, the system delivers ∼93 dB sensitivity with approximately 1.5 mW of sample illumination power (see Methods). An integrated epi-fluorescence and bright-field module, implemented via a dichroic mirror placed between the tube lens and the fold mirror, allows simultaneous acquisition of widefield images alongside OCM volumes. Data acquisition, galvo synchronization, and hardware triggering are orchestrated by a custom LabVIEW program via an FPGA board, while raw spectral data are streamed to solid-state drives or transferred to a GPU for real-time processing and visualization. Critically, the entire optical assembly was miniaturized into a compact module measuring 52.5 × 42.5 × 13 cm with a total system weight of approximately 25 kg, enabling transport in a standard heavy-duty suitcase and deployment at remote field sites beyond conventional laboratory infrastructure (**Fig. 1b-d, SI Fig. 1**).

**Figure 1.**
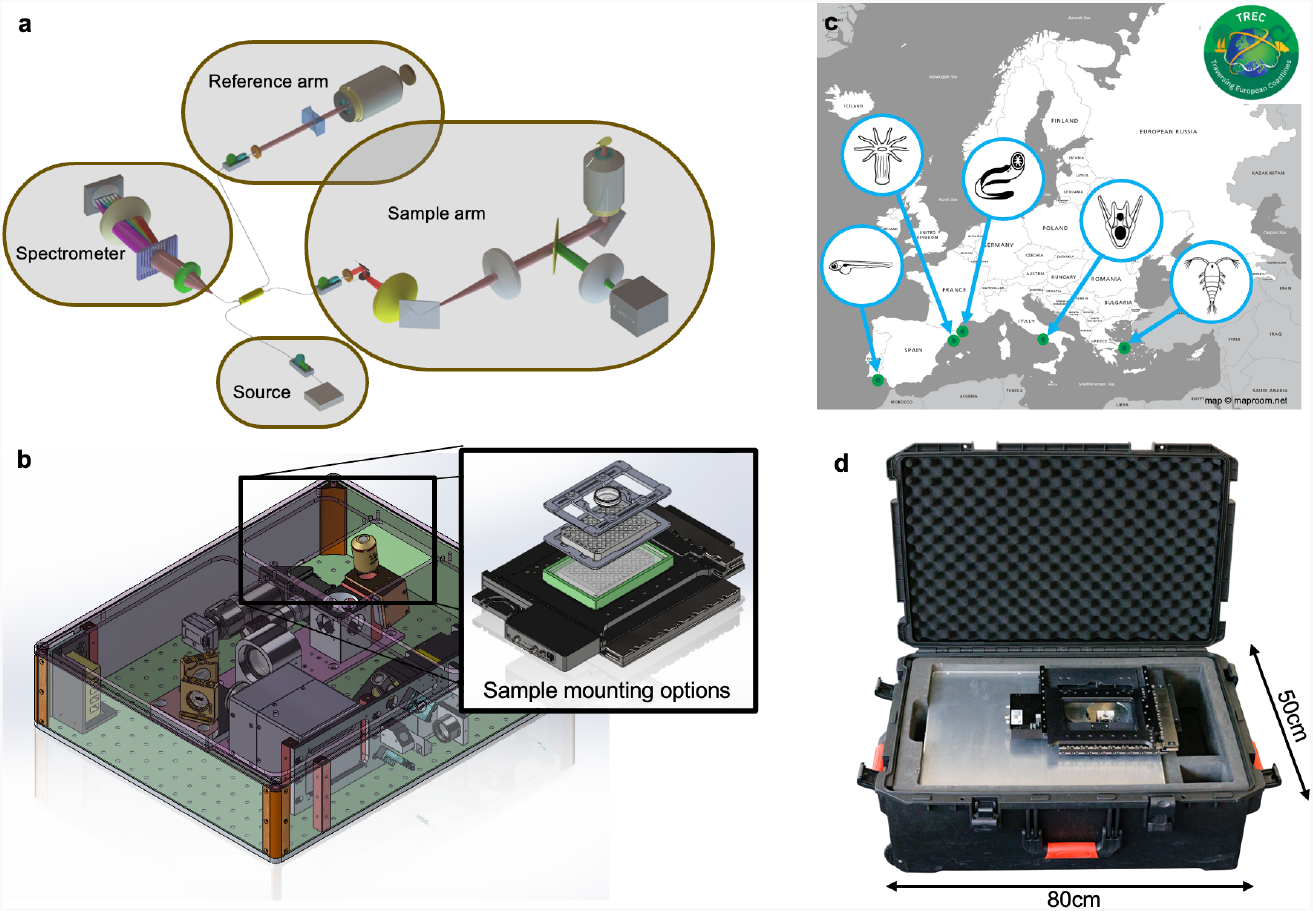
Mobile OCM system design and field deployment. **(a)** Schematic diagram of the spectral-domain OCM system, illustrating the fiber coupler-based Michelson interferometer with its three main components: the broadband SLD light source, the reference arm with dispersion-compensating optics, and the sample arm incorporating a galvo mirror pair in a 4f scanning configuration with interchangeable objective lenses. Backscattered interference light is directed to a custom high-speed spectrometer (also see **SI Fig. 1**). **(b)** Left: CAD rendering of the compact optical assembly, showing all components integrated into a 42.5 × 52.5 × 13 cm enclosure mounted on a standard optical breadboard. Right: modular sample mounting options, including a multiwell plate adapter, stage-top incubator and standard coverslip stage, enabling flexible sample configurations for different organism sizes and imaging geometries. **(c)** Map of European field deployment sites including stops of the EMBL TREC expedition. Green dots indicate sampling and OCM imaging locations (L-R: Faro, Barcelona, Banyuls-sur-Mer, Naples, Athens) with exemplary aquatic invertebrate taxa imaged at each site indicated by icons. **(d)** The fully assembled mobile OCM system packed in a standard heavy-duty transport suitcase, illustrating the field-deployable form factor enabling operation at remote sites beyond conventional laboratory infrastructure.

### Cross-phyla structural imaging of aquatic metazoans

To demonstrate the versatility of the mobile OCM platform, we imaged a broad range of aquatic invertebrates both in the lab as well as in the field, spanning multiple phyla, including cnidarians, poriferans, annelids, and arthropods, as well as the vertebrate model *Danio rerio* (**Fig. 2**). Maximum-intensity projections across all specimens confirm that the intrinsic scattering contrast of OCM is broadly applicable to soft-bodied aquatic organisms, resolving internal anatomy and tissue boundaries across millimeter-scale volumes without fluorescent labeling.

**Figure 2.**
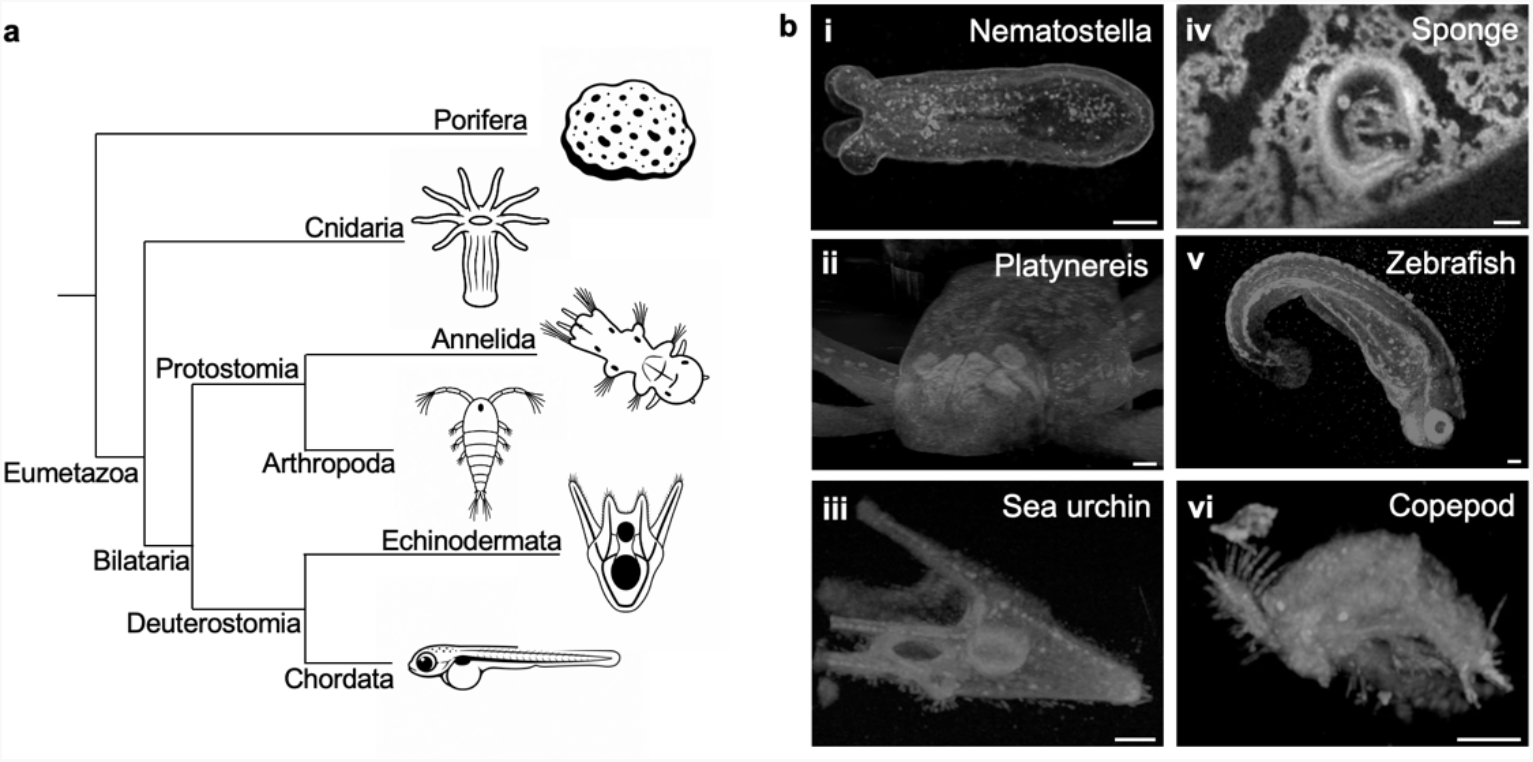
Cross-phyla overview of aquatic invertebrates imaged by mobile OCM. **(a)** Schematic phylogenetic overview of the aquatic invertebrate phyla represented in this study (Cnidaria, Annelida, Arthropoda, Echinodermata, Porifera) alongside the vertebrate model *Danio rerio*, with representative line drawings illustrating body plans. **(b)** Representative maximum-intensity projection OCM images of each organism. **(i)** *Nematostella vectensis* (Cnidaria), showing the elongated body column and internal gastrovascular cavity. **(ii)** *Platynereis dumerilii* (Annelida), showing an anterior view of the dissected head of a premature worm. **(iii)** Sea urchin tube feet (Echinodermata). **(iv)** Freshwater sponge (*Spongilla lacustris*, Porifera), showing the aquiferous internal canal system and a prominent gemmule (asexual reproductive cyst). **(v)** Zebrafish larva (*Danio rerio*), showing the curved body axis, notochord, and head structures including the eye. **(vi)** Copepod (Arthropoda), resolving the cephalothorax, articulated appendages, and fine surface structures. Scale bars: 100µm, 10µm for (iii).

We next examine the structural imaging capabilities in detail for four representative organisms, using orthogonal cross-sectional views to highlight phylum-specific anatomical features (**Fig. 3**). More specifically, in Hydra, OCM imaging revealed the layered organization of the body column in both longitudinal and transverse views (**Fig. 3a**). The side-view projection shows the part of the body column including the aboral pole, while the en face cross-section resolves the tubular body wall as a distinct annular structure enclosing the central gastrovascular cavity, demonstrating that OCM can resolve the characteristic two-layered body plan of this cnidarian at micrometer-scale resolution without any exogenous labeling.

**Figure 3.**
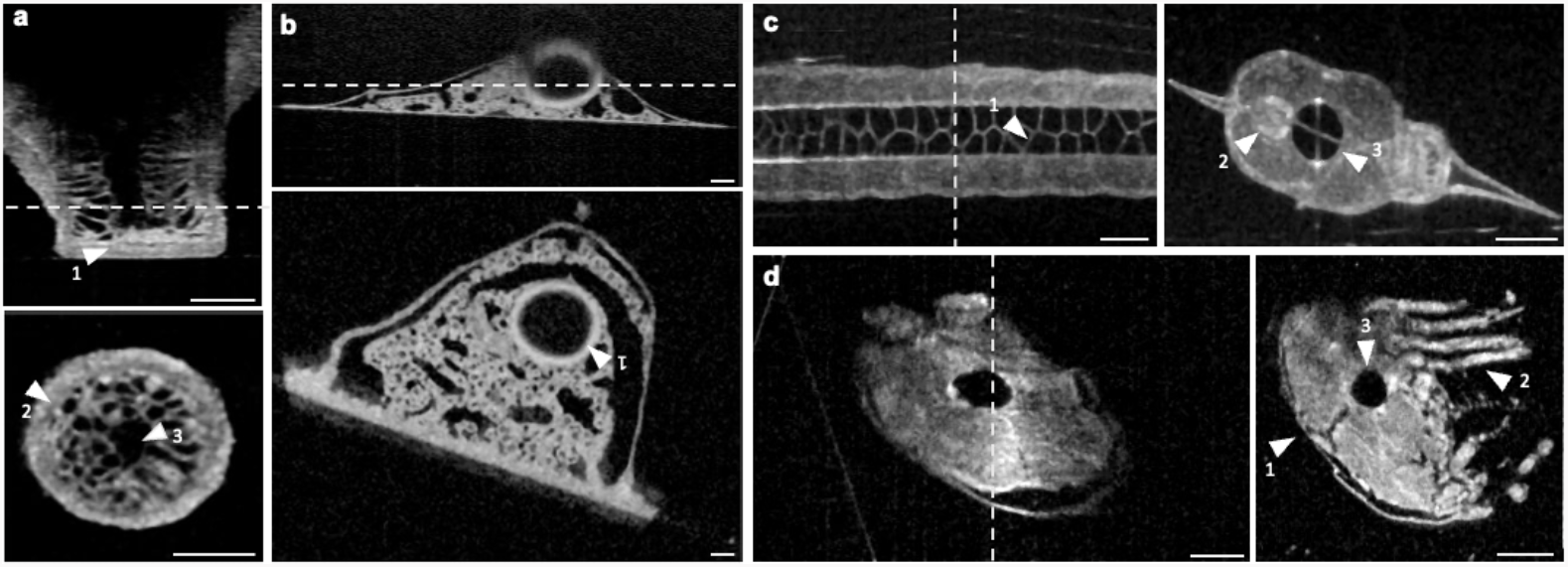
Cross-phyla structural imaging of aquatic invertebrates by mobile OCM. Orthogonal OCM image views of four aquatic organisms demonstrating label-free, three-dimensional structural imaging across phyla. **(a)** Hydra (*Cnideria*): longitudinal side-view projection (top) and en face cross-section at the depth indicated by the dashed line (bottom), resolving the two-layered tubular body wall enclosing the central gastrovascular cavity. Arrowheads indicate: 1, basal disc; 2, ectodermal-endodermal tissue boundary; 3, gastrovascular cavity lumen. **(b)** Freshwater sponge (*Spongilla lacustris*): longitudinal view (top) and en face cross-section (bottom) revealing the interconnected aquiferous canal system. Arrowhead (1) indicates a gemmule (asexual reproductive bud); epithelial-like pinacocytes covering the sponge and choanocyte chamber are visible throughout. **(c)** Zebrafish larva (*Danio rerio*): longitudinal view through the trunk (top left) and transverse cross-section (top right) resolving individual vacuoles (1), as well as the spinal cord (2) and notochord (3). **(d)** Copepod (*Arthropoda*): longitudinal view (left) and transverse cross-section (right) resolving the prosome^24^ (1), thoracic legs^25^ (2) and internal body structures including putative digestive organs (3). All scale bars: 100 µm.

**Fig. 3b** shows label-free OCM images of freshwater sponge (*Spongilla lacustris*) and exemplifies detailed visualization of its characteristic aquiferous canal system, water pumping choanoderm with choanocyte chambers, the epithelial-like pinacoderm ‘tent’ and internal mesohyl tissue in three dimensions. Cross-sectional and en face views reveal a complex network of incurrent and excurrent canals and choanocyte digestive chambers permeating the sponge, demonstrating that OCM can image the sponge’s internal architecture at near-cellular resolution over hours or days, revealing the detailed anatomy, physiology, and behaviors in this nerveless animal^22^.

In zebrafish larvae, OCM demonstrated the system’s capability for resolving fine internal anatomy in a vertebrate model organism (**Fig. 3c**). A longitudinal cross-section through the trunk region reveals the characteristic layered tissue organization, including key anatomical landmarks such as the notochord and surrounding somitic muscle blocks. The ability to resolve these features label-free in a live fish larva illustrates the broad applicability of OCM contrast beyond invertebrate specimens.

As another example application of high-resolution OCM, we imaged small crustaceans with complex surface geometries. Specifically, OCM images of a copepod revealed the detailed three-dimensional anatomy of this small crustacean under near-native conditions (**Fig. 3d**). The volumetric data resolves the copepod’s body organization and underscores the capability of mobile OCM to image morphologically complex, semi-transparent organisms at sufficient resolution to distinguish internal soft-tissue compartments without the need for mounting, fixation, or contrast agents. By resolving distinct morphological variations in specimens collected from diverse habitats, OCM can be a valuable tool for field expeditions and biodiversity assessments.

### Morphodynamic imaging

*nematosome tracking, cell division, and peristaltic gut motility* Beyond static structural characterization, the high imaging speed of the mobile OCM enables time-resolved volumetric imaging of dynamic biological processes across a range of aquatic organisms. To demonstrate this capability, we captured three distinct classes of morphodynamic events spanning cellular aggregate motility, cell division, and organ-level tissue dynamics. In *Nematostella vectensis*, we used fast time-lapse OCM to track the movement of nematosomes^26^ - motile cellular aggregates that patrol the gastrovascular cavity and might function as immune-like cell clusters^27^ - at the volume rate of 1.85 Hz. OCM’s intrinsic scattering contrast resolves nematosomes as discrete high-contrast objects against the surrounding lumen, enabling particle tracking over extended time periods without any labeling. Overlaying color-coded speed tracks onto the OCM volume projection (**Fig. 4a**) reveals the characteristic active, directed trajectories of individual nematosomes traversing the body column at speeds ranging from ∼0.1 to ∼100 µm/s. This confirms that OCM can resolve and track these structures with sufficient spatial and temporal resolution to quantify their motility parameters in a living, intact animal, which would be impossible from 2D images only. This capability is opening the door to probing nematosome immune-like function by quantifying motility responses upon pathogen challenge or immune stimulation.

**Figure 4.**
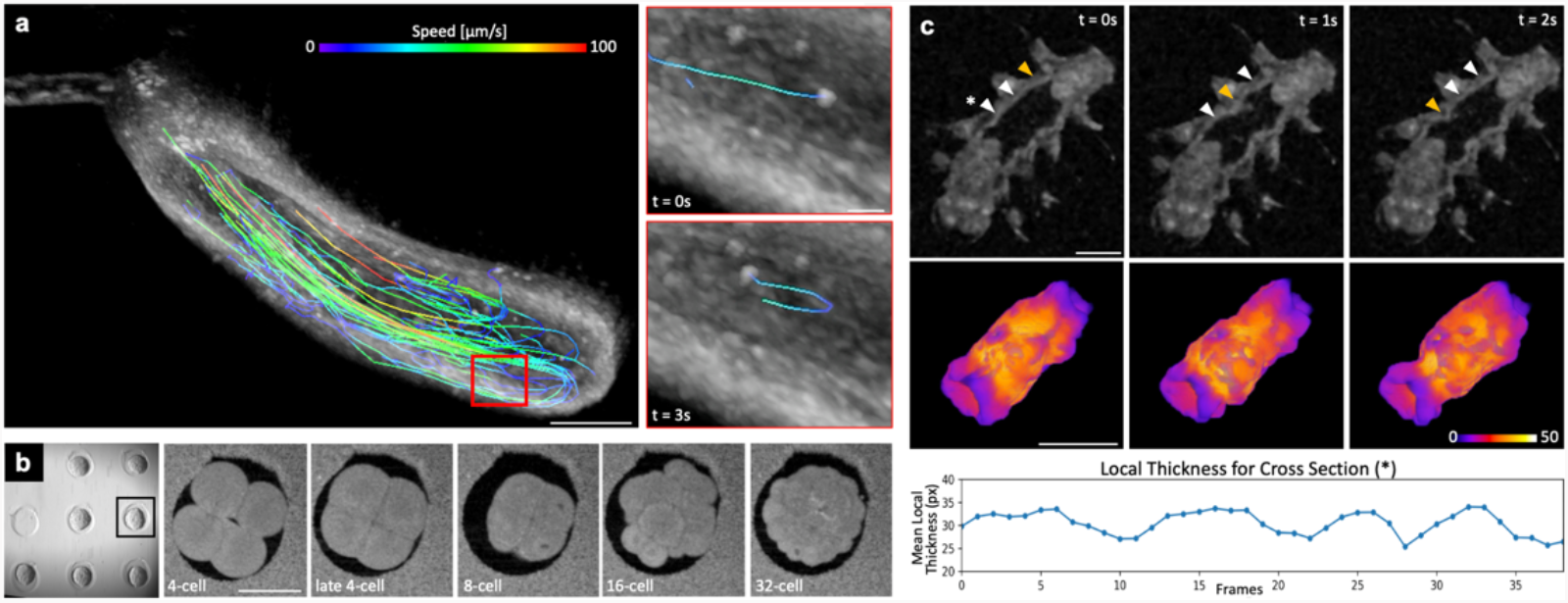
Morphodynamic imaging of aquatic invertebrates by mobile OCM. **(a)** Nematosome tracking in *Nematostella vectensis* (Cnidaria). Left: maximum-intensity projection of an OCM volume with color-coded particle tracks overlaid, showing nematosome trajectories color-mapped by instantaneous speed (0–100 µm/s). The red box indicates the region enlarged in the insets. Right insets: close-ups of a single tracked nematosome at t = 0 s and t = 3 s, illustrating its spatial displacement along the trajectory. Scale bar (left): 100 µm; inset scale bar: 20 µm. See **Supplementary Video 1. (b)** Early embryonic cell divisions in ascidian embryos. Left: overview of a multiwell plate showing multiple embryos; boxed embryo is shown in the time series at right. Sequential en face OCM frames capture stereotyped cleavage progression from the 4-cell stage through late 4-cell, 8-cell, 16-cell, and 32-cell stages over approximately 2 hours. Scale bar: 100 µm. See **Supplementary Video 2. (c)** Peristaltic gut motility in starved *Platynereis dumerilii (*Annelida). Top row: Longitudinal views at t = 0, 1, and 2 s showing a z-slice of the whole organism with its gut lumen and surrounding tissue. Orange arrowheads indicate a region of active tissue contraction; white arrowheads track a discrete tissue feature undergoing displacement consistent with directed peristaltic transport. See **Supplementary Video 3**. Bottom row: 3D segmentation of gut, color coded based on the local thickness. Graph displays the mean local thickness across a transverse cross section (*), highlighting the rhythmic contraction over time. Scale bar: 100 µm. One frame equals 0.263 sec. Also see **Supplementary Video 4**.

We further demonstrated the high-throughput capability of the mobile OCM and its compatibility with standard multiwell dish formats by imaging ascidian embryos undergoing early cleavage divisions (**Fig. 4b**). In particular, OCM was able to capture the stereotyped progression of symmetric cell divisions, and the transition from a multi-lobed early cleavage stage through progressive rounds of division toward a more compact, rounder morphology. The ability to image embryos directly in standard tissue culture vessels without specialized mounting underscores the practical accessibility of the platform for high-throughput and parallelized developmental biology applications, including time-lapse studies of embryogenesis in non-model organisms.

Finally, we imaged peristaltic gut motility in *Platynereis dumerilii* over three consecutive timepoints (**Fig. 4c**). Paired cross-sectional (top row) and longitudinal (bottom row) views reveal the dynamic repositioning of gut contents during peristaltic movement imaged a 3.8Hz volume rate. The circular gut lumen and surrounding tissue wall as well as their peristaltic contractile activity could be well discerned, demonstrating that OCM can capture organ-level tissue dynamics in a live annelid at sufficient temporal and spatial resolution to distinguish intrinsic tissue motion, entirely without fluorescent labeling or tissue clearing.

### Imaging of marine plankton and field-collected specimens

Beyond established laboratory model organisms, we further demonstrated the capability of the mobile OCM for rapid, label-free structural imaging of marine plankton collected during the EMBL Traversing European Coastlines expedition (TREC) expedition led by EMBL, in collaboration with EMBRC and the Tara Oceans Foundation, which took place between 2023-2024. Over 18 months, researchers collected comprehensive samples across 21 countries to investigate critical ecological issues, including biodiversity loss, pollution, antibiotic resistance, and the anthropogenic impact on habitats (https://www.embl.org/about/info/trec/). Figure 5a shows a representative brightfield overview of a mixed plankton sample, illustrating the morphological diversity of co-occurring organisms captured from a single field collection. OCM imaging of individual organisms from this community revealed taxon-specific structural features across a wide range of body plans and size scales, without any fixation, staining, or specialized sample preparation.

**Figure 5.**
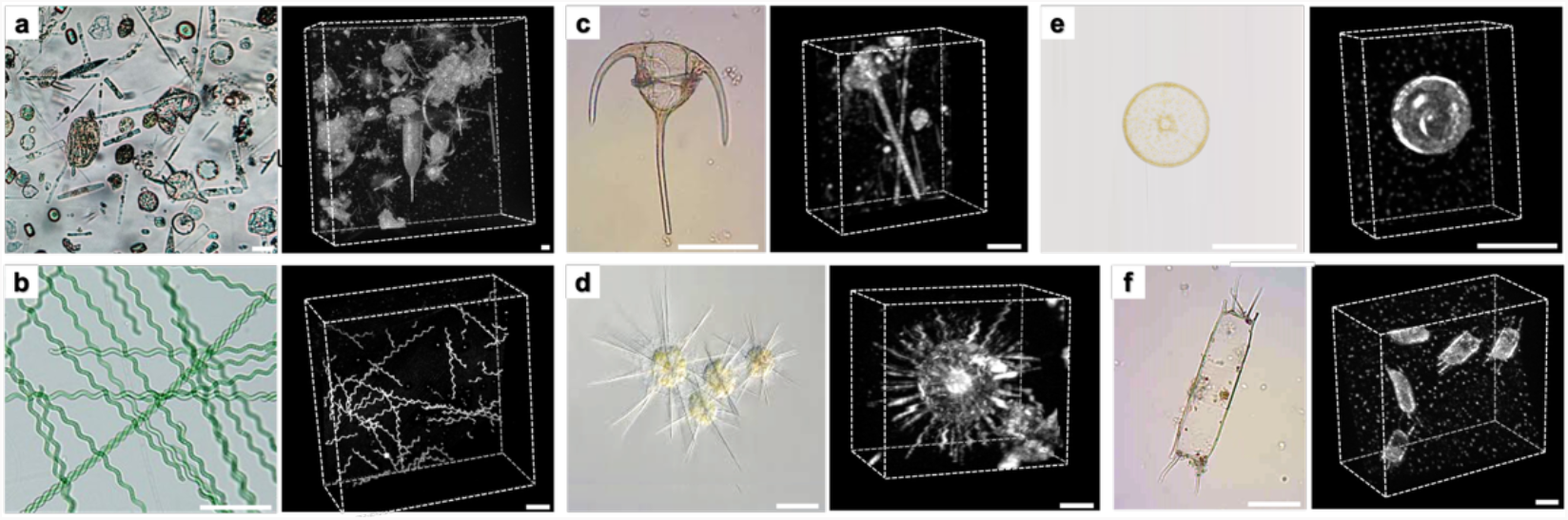
Label-free 3D phenotyping of marine plankton by mobile OCM. **(a)** Brightfield overview of a mixed marine plankton sample collected during the EMBL TREC expedition, illustrating the morphological diversity of co-occurring organisms from field collections. **(b-f)** For each taxon, an exemplary brightfield reference image (left) and label-free, volumetric OCM image (right) are shown side by side. **(b)** *Arthrospira* sp. (Cyanobacteria), with its characteristic helical filament geometry. **(c)** *Ceratium tripos* (Dinoflagellata), showing the triradiate horn morphology and internal cell body. **(d)** *Acantharia* (Radiolaria), revealing the ornate spine arrangement and central capsule. **(e)** *Coscinodiscus granii* (Bacillariophyta), rendered as a well-defined circular frustule with visible surface patterning consistent with its elaborately structured silica cell wall. **(f)** *Odontella sinensis* (Bacillariophyta), with its elongated silica frustule and showing internal chamber organization. Scale bar: 100µm, 50µm for (C).

In the filamentous cyanobacterium *Arthrospira*, OCM resolved the characteristic helical filament geometry with high contrast against the background medium, faithfully reproducing the regular pitch and diameter of the trichomes visible in brightfield (**Fig. 5b**). In the dinoflagellate *Ceratium tripos*, the triradiate horn morphology and internal cell body were clearly delineated in three dimensions, with OCM providing complementary contrast to brightfield by rendering internal structural boundaries that are poorly visible in transmission (**Fig. 5c**). The siliceous skeleton of the radiolarian *Acantharia* was also clearly rendered and its elaborate spine arrangement and central capsule geometry resolved in detail consistent with the organism’s ornate mineralized architecture (**Fig. 5d**). In the centric diatom *Coscinodiscus granii*, the circular frustule was rendered as a well-defined disc with visible surface patterning, consistent with its elaborately structured silica cell wall (**Fig. 5e**). Finally, for the pennate diatom *Odontella sinensis*, OCM captured the elongated frustule outline and internal chamber structure, with the silica cell wall appearing as a high-scattering shell enclosing the lower-scattering cytoplasm (**Fig. 5f**).

Taken together, these results establish that mobile OCM can deliver high-resolution, three-dimensional structural imaging of morphologically diverse marine plankton under near-native, field-compatible conditions, also of smaller sized, ∼100µm large specimens. The ability to pair OCM directly with brightfield (fluorescence) overview imaging of bulk plankton samples, without any intermediate preparation steps, illustrates a practical workflow for rapid label-free phenotyping of natural plankton communities, with potential applications in biodiversity monitoring and environmental assessment at remote sampling sites.

## Discussion and Outlook

Our mobile OCM addresses a longstanding disconnect between the resolution demands of modern cell and developmental biology and the practical constraints of studying organisms outside the laboratory. By bringing micrometer-scale, label-free, volumetric imaging to the field, it enables a class of experiments that was previously impossible: characterizing the morphology and dynamics of fragile marine organisms across phyla immediately upon collection, before transport-induced stress, behavioral alteration, or organismal death obscures the biology of interest. The TREC expedition deployments reported here represent, to our knowledge, the first application of high-resolution OCM to field-collected marine organisms and plankton imaged on-site, and illustrate a practical path toward integrating advanced optical microscopy into environmental research workflows^5^. Here, in the future additional environmental perturbations could be applied to the specimens under investigations, and associated changes quantitively be recorded over space and time. More broadly, the cross-phyla demonstrations spanning cellular aggregate tracking in cnidarians, embryonic cleavage in ascidians, and organ-level peristalsis in annelids, establish that label-free OCM contrast is sufficiently general and informative to support quantitative morphodynamic studies across the full diversity of aquatic animal life, including unicellular eukaryotes and organisms where fluorescent transgenesis remains technically challenging or undesirable.

A key advantage of OCM for biological imaging of aquatic organisms is its use of near-infrared illumination, which is of low phototoxicity and minimally perturbs native, living samples, an important consideration given that visible light employed in light-sheet and confocal microscopy can induce stress responses and alter the very biological processes under observation. By imaging biological specimens under near-native conditions and without immobilization, chemical fixation or labeling, OCM can capture morphological and dynamic information that more closely reflects the organism’s physiological state in its natural environment. This is particularly valuable for studying slow contractile processes, developmental transitions, and tissue dynamics in organisms where fluorescent transgenesis remains technically challenging or undesirable. Furthermore, the interferometric detection of the backscattered light makes OCM intrinsically immune to background light so that the OCM can also be operated in bright daylight conditions as encountered outside the laboratory, unlike many traditional (fluorescence) imaging techniques.

Several technical avenues offer opportunities to further expand the capabilities of the platform. The current axial resolution of ∼2.5 µm in water is determined by the effective spectral bandwidth of the SLD source and the system’s spectral response function. Broader bandwidth sources, or the use of visible-range OCM, could push axial resolution toward or below 1 µm, enabling sub-cellular discriminability^28,29^. We note that lateral resolution, which degrades with defocus at higher numerical apertures, remains a fundamental challenge for imaging thick specimens: future implementations could incorporate non-diffractive (Bessel-beam) beam engineering and/or computational refocusing approaches to recover diffraction-limited lateral resolution throughout extended depth ranges^30^. Similarly, transition to longer wavelength ranges would reduce scattering in opaque tissues and thereby extend penetration depth at the cost of more complex hardware^31^. Full-field OCM configurations, which illuminate and detect in parallel rather than point-by-point, represent another avenue for increasing volumetric imaging throughput and sensitivity, potentially enabling imaging at cellular resolution over much larger fields of view^32^.

The field deployments conducted during the EMBL TREC expedition demonstrated that the mobile OCM can be reliably operated at remote sites with minimal setup overhead. This positions OCM as a natural complement to the growing ecosystem of advanced mobile laboratory infrastructure^5^ that brings high-end analytical capacity directly to the sampling site. By minimizing the delay between sample collection and imaging, and thus the impact on the organism, mobile OCM opens new avenues for studying non-model organisms and complex environmental communities in their native context. Here, the mobile OCM is ideally positioned to integrate into such platforms: its compact footprint, suitcase-transportable form factor, and label-free contrast mechanism make it directly compatible with the operational philosophy of mobile labs which aim to provide rapid, minimally invasive (pre-)analysis of live organisms immediately post-collection. Specifically, OCM fills a complementary niche by providing fast, penetrating, three-dimensional structural imaging of larger, optically dense specimens, including intact invertebrates, tissue fragments, and macroscopic marine organisms that are poorly served by confocal or light-sheet approaches. In this respect, a possible future application and workflow is the combination of OCM with active sorting or fixation based on morphology and/or morphodynamics.

More broadly, mobile OCM exemplifies a shift in biological imaging toward instrumentation that follows the biology as the scientific community increasingly recognizes the importance of studying organisms in, or close to, their native habitats, particularly in the context of climate change, biodiversity loss, and environmental monitoring. For these, field-deployable, non-invasive imaging modalities will play an essential role. Future implementations of mobile OCM could incorporate automated scanning, microfluidics and image analysis pipelines for high-throughput morphometric biodiversity assessment, followed by active sorting and complementary post-imaging -omics characterization. Together with advanced mobile laboratory platforms and, mobile OCM has the potential to establish a new paradigm for integrative, field-ready 3D bioimaging, enabling quantitative structural and dynamic studies of aquatic life across evolutionary scales and environmental contexts.

## Methods

### Spectral-domain Optical Coherence Microscopy

*Setup:* The near-infrared light from a superluminescent diode (SLD) is directed into the source arm of a 20/80 fiber coupler-based Michelson interferometer. The light is split into the reference and sample arms, with the larger portion directed into the reference arm and the smaller portion into the sample arm. Two identical *f*=12 mm collimator lenses (60FC-4-M12NIR-10, Schäfter + Kirchhoff GmbH, Hamburg, Germany) at the exits of the fiber coupler direct and collimate the light in the free-space sections of the reference and sample arms. Polarization is matched by manually tuning the fiber polarization controllers. Two types of SLDs (EBD312018-31, Exalos AG, Schlieren, Switzerland, and M-T-850-HP-I, Superlum Diodes Ltd, Cork, Ireland) were used in our experiments, both with similar 3dB bandwidths of ∼160 nm, which resulted in a theoretical resolution of ∼2.6 μm in air (∼2.0 μm in water).

In the sample arm of the OCM, the collimated light was scanned by a galvo mirror pair (P5XA101A, Scanner Optics Co., Ltd. Shenzhen, China), which was conjugated to the back focal plane of the objective (CFI Plan Achro 2× NA 0.06, Plan Fluor 4× NA 0.13, and Plan Fluor 10× NA 0.30, Nikon, Japan) to form a ‘4f’ configuration with the scanning lens and the tube lens (LSM54-850 and TTL200MP, Thorlabs, Newton, USA). A fold mirror was placed between the tube lens and the objective to reflect the light beam upwards at a right angle. To minimize specular reflectance from the coverslip surface, both the fold mirror and the objective lens were slightly tilted by 8° relative to normal incidence.

The reference arm was composed by a focusing lens, a retroreflector, five reflective mirrors and a dispersion compensator. The retroreflector was mounted and aligned on a translation stage to ensure that the optical path lengths in the reference and sample arms were matched. The optical dispersion introduced by the different optical elements in both arms was compensated by a carefully adjusted prism pair together with other dispersion compensators.

The back-scattered signal from the sample and the reference light reflected from the mirror interfered at the fiber coupler and were subsequently directed to a custom-built spectrometer through the fiber in the detection arm. The line camera used in the spectrometer (Octoplus, Teledyne e2V, Saint-Egrève, France) has a maximum read-out speed of 250 kHz. The spectrometer was designed to cover a wavelength range from 760 nm to 920 nm.

#### Performance

The maximum imaging speed of the OCM system is 250k A-lines/s, which is constrained by the speed of the spectrometer camera. In our experiments, we achieved a maximum volume rate of 7.7 Hz (180×180 pixels laterally). The light power illuminating the samples was approximately 1.5 mW (slightly adjusted for different objective lenses), resulting in a measured sensitivity of ∼93 dB with an exposure time of 4 µs. The actual axial resolution was 3.35 µm in air, corresponding to 2.52 µm in water, as determined by fitting the PSF of the signal peak from a mirror placed near zero delay. This resolution was lower than the theoretical value predicted by the light source’s spectral bandwidth of 160 nm. The reduced effective bandwidth is attributed to the apodization of the recorded spectra by the spectral response function of the entire experimental system. The theoretical lateral resolutions at the focal plane are 7.0 µm, 3.57 µm, and 1.44 µm for the 2×, 4×, and 10× objective lenses, respectively. It is important to note that the lateral resolution deteriorates when out of focus, with higher magnification objective lenses experiencing faster degradation. In contrast, the axial resolution slightly degrades with increasing optical delay, as it depends on the system’s spectral resolution and bandwidth^33^.

### Epi-fluorescence and bright field microscopy

To simultaneously capture regular widefield images of the sample, a dichroic mirror with a cutoff wavelength of 662 nm (F38-662_T3, AHF Analysentechnik AG, Tübingen, Germany) was positioned between the tube lens and the flip mirror to reflect visible light. Depending on the required magnification, an achromatic lens with a specific focal length (e.g., *f* = 100 mm or 200 mm, AC254-100-A, AC254-200-A, Thorlabs, Newton, USA) was selected to focus the light onto a CMOS camera (CM3-U3-50S5M-CS, Teledyne FLIR, Frankfurt, Germany). In epifluorescence mode, a filter cube containing a dichroic mirror, excitation filter, emission filter, and an LED light source was used. The wavelength specifications for these optical elements were chosen based on the fluorochrome of the sample. The epi-fluorescence modality could be switched to bright field by removing the filters from the filter cube. For bright field imaging, the illumination could either be ambient light or a halogen cold light source directed at the sample.

### Data acquisition

The interference spectrograms recorded by the camera were transferred to the workstation memory via camera link cables and a frame grabber board (AXN-PC2-CL-1xE, BitFlow, Woburn, USA). The raw spectral data in memory were either streamed directly to a solid-state drive (SSD) for storage or copied to the CPU or GPU (graphics processing unit) for processing and visualization. When fluorescence or bright-field images were acquired simultaneously, they were streamed to a separate SSD. The workstation was equipped with an NVIDIA GeForce GTX 1080 GPU (PCIe 3.0, 2560 cores at 1.6 GHz, 8 GB GDDR5X graphics memory). Image acquisition and recording were controlled by a custom LabVIEW program, which synchronized the galvo mirror pair, the spectrometer, and other hardware components via an FPGA board (PCIe-7841R, National Instruments, Austin, USA). The laser beams fast scanning direction, driven by the galvo mirrors, could follow either a raster or zigzag pattern. In most of our experiments, the fast scanning followed a zigzag pattern during image acquisition. The magnifications along the X and Y axes of the microscope were calibrated in advance using a CMOS camera with a large sensor.

### Data processing pipeline

The depth scan profiles (A-lines) were reconstructed from the interference spectral signal using a standard OCT postprocessing procedure^33^. This included wavenumber linearization, background subtraction, spectrum reshaping, and inverse fast Fourier transformation (IFFT). If significant residual optical dispersion remained, digital dispersion compensation was applied after spectrum reshaping^34^. Finally, the intensity of the resulting image was scaled using a logarithmic grayscale.

### Sample preparation, imaging and postprocessing

#### Nematostella

For imaging, live non-anesthetized polyps were mounted in a single drop of 12 ppt artificial sea water (sea salt, instant ocean) on a glass bottom dish (MatTek no. P35G-1.5-14-C), with a second coverslip on top. The polyps were anesthetized with 7% MgCl2 and imaging was performed with a 4x objective lens (Nikon, Plan Fluor, NA 0.13).

#### Spongilla lacustris

Gemmules (protected packets of dormant stem cells that function as asexual buds and ‘hibernation’ strategy) of S. lacustris embedded in previous years sponge remains were collected on 22 February 2023 from Lake Constance, near Kressbronn, Germany (47°35’09.0”N 9°35’56.4”E). The gemmules were extracted from adult tissue by gently rubbing the sponge patches over grit 500 sandpaper and stored at 4°C in water. To culture sponges for OCM imaging, 2-4 gemmules were placed in glass-bottom culture dish (Greiner Bio-One cat# 627860) containing 5mL sterile filtered water from Lake Constance and kept at 18°C in the dark. After one week of growth, the sponges adhered to the glass surface and were ready for imaging.

For live sponge OCM imaging, individual 8-day old Spongilla specimens were grown in 35 mm glass-bottom culture dishes (Greiner Bio-One cat# 627860) containing 5 mL of filtered lake water. Natural attachment of the Spongilla specimen to the glass bottom enabled consistent imaging using the inverted geometry. The speed of the OCM was set to ∼5 s/volume (exposure time 20 µs for each A-line) resulting in higher signal-to-noise ratio and image contrast, and we chose a 4x objective lens (NA=0.13) which achieved near-isotropic spatial optical resolution of ∼2.5 µm. Each volume scan contained 512×512 steps along x and y direction with a step size of 5.05 µm, resulting in a field of view of 2.6 mm. We note that the spatial laser scanning step resolution was approximately twice as large as the optical resolution, which we found sufficient for high-quality segmentation, but kept overall data size of the longitudinal recordings manageable.

#### Platynereis dumerilii

Animals were raised at 18°C in FNSW (filtered natural seawater) with a 16h light: 8h dark cycle and under an artificial moon cycle (3 weeks of darkness, 1 week of artificial moonlight). A premature worm (>50 segments) was anaesthetized (1:1 mixture of 7.5% MgCl and FNSW) and its head was dissected above the pharynx on ice and placed on a glass-bottom culture dish for OCM imaging (Fig. 2b). For live-imaging of gut motility (Fig. 4c), a juvenile worm (5-segments) was starved for 72h and then placed on a glass-bottom culture dish for OCM imaging.

OCM images of *Platynereis* were segmented into foreground (tissue) and background using pixel classification in Ilastik^35^ (v1.4.0-OSX), after which the gut lumen was extracted using a custom processing script in FIJI^36^. Local thickness of the gut lumen was computed using the localthickness python package^37^.

#### Zebrafish

The AB2B2 zebrafish (*Danio rerio*) wild-type line was used in this study. All zebrafish experiments were conducted in accordance with the EMBL Institutional Animal Care and Use Committee (IACUC code: 22-003_HD_MD). The larvae used in this study were raised from 72 hours post-fertilisation (hpf) at 28.5°C. At 6 hours post-fertilisation (hpf), some of the larvae were transferred to 36°C. Adult fish were kept at 28.5°C under light/dark cycles of 14–10 hours and fed daily with hatched Artemia. Zebrafish embryos were maintained in Petri dishes containing 1x E3 solution (for the 60x stock solution: 17.2 g NaCl, 0.76 g KCl, 2.9 g CaCl_2_·2H_2_O, 4.9 g MgSO_4_·7H_2_O, 1 L distilled H_2_O). At 24 hpf, we replaced the E3 solution with one containing 0.003% w/v N-phenylthiourea (PTU, Sigma-Aldrich, 103-85-5) to prevent pigmentation in the developing larvae. The larvae were imaged while still alive. Prior to imaging, the zebrafish larvae were anaesthetised using MESAB (ethyl 3-aminobenzonate methanesulfonate salt, Sigma-Aldrich, A5040-100G). We used 35 mm glass-bottom dishes for imaging, adding the fish with fresh PTU and MESAB.

#### Hydra

Specimens of *Hydra vulgaris* AEP were starved for 24h and deposited over the glass bottom of a water-tight dish (µ-Dish, Ibidi, Cat.No:81156), where the animal would attach after 10-15 min. Finally, the dish was flipped upside-down to expose the basal disc and lower peduncle regions for imaging.

#### Environmental plankton sample collection

Plankton samples were collected by towing nets of various sizes (20-150 µm) for 2-5 minutes, depending on the site. Once recovered on the boat, the samples were further fractionated using metal sieves (500 µm or 200 µm), transferred into bottles and kept at ambient seawater temperature in a cooler. Upon arrival, samples were kept in an incubator at ambient seawater temperature running a 12 hours day/night cycle and processed within 4 hours of retrieval. When necessary, cells were gently concentrated onto a 1.2 µm mixed cellulose ester (Millipore) filter using a manual vacuum filtration unit. Once the volume was sufficiently reduced, cells were gently resuspended from the filter and put onto glass bottom dishes for OCM imaging.

## Acknowledgements

We would like to acknowledge support by the EMBL Heidelberg mechanical and electronic workshops and the Mobile Laboratories Core Facility. We thank the TREC Consortium members, TREC Expedition team and TREC local partners for permits, hosting and guidance on sampling sites. We thank the Institut de Ciències del Mar (ICM-CSIC) Spain, the Stazione Zoologica Anton Dohrn, Italy and the Hellenic Centre for Marine Research, Greece for their hospitality and support during the TREC expedition. We are further grateful to Madison Bolger-Munro and Heisenberg group (IST Austria) for providing the ascidian samples shown in Fig. 4b, and to Johan Decelle (Université Grenoble Alpes) for help with radiolarian samples. J.M. acknowledges support by a Research Grant from HFSP (Ref.-No:1936 RGP024/2023). R.P. acknowledges support of an ERC Consolidator Grant (no. 864027, Brillouin4Life), and the Horizon Europe Framework Programme (101094250/IMAGINE). This work was supported by funds from the European Molecular Biology Laboratory.

## Disclosures

The authors declare no competing interests.

## Data availability

The raw data underlying the results presented in this paper are not publicly available at this time but will be made public at time of publication.

## Author contributions

L.W. conceptualized, designed and built the mobile OCM platform. L.W. performed the imaging experiments with assistance from S.D., R.M., A.S., F.R., and T.Q.. L.W. and A.S. analyzed the data, images, and videos and generated the figures together with S.D. and T.Q.. T.B., M.B., M.L., R.M., F.R., T.W., V.A.W. contributed samples and assisted in data interpretation, under the supervision of M.D., J.S., J.M., D.A., N.L., F.V., and A.I.. L.W. and A.G.O. designed and built custom electronic components. R.P. conceived and supervised the project, acquired funding, and wrote the manuscript, with input from L.W. and all other authors.

## Supplementary Figures

**Supplementary Figure 1.**
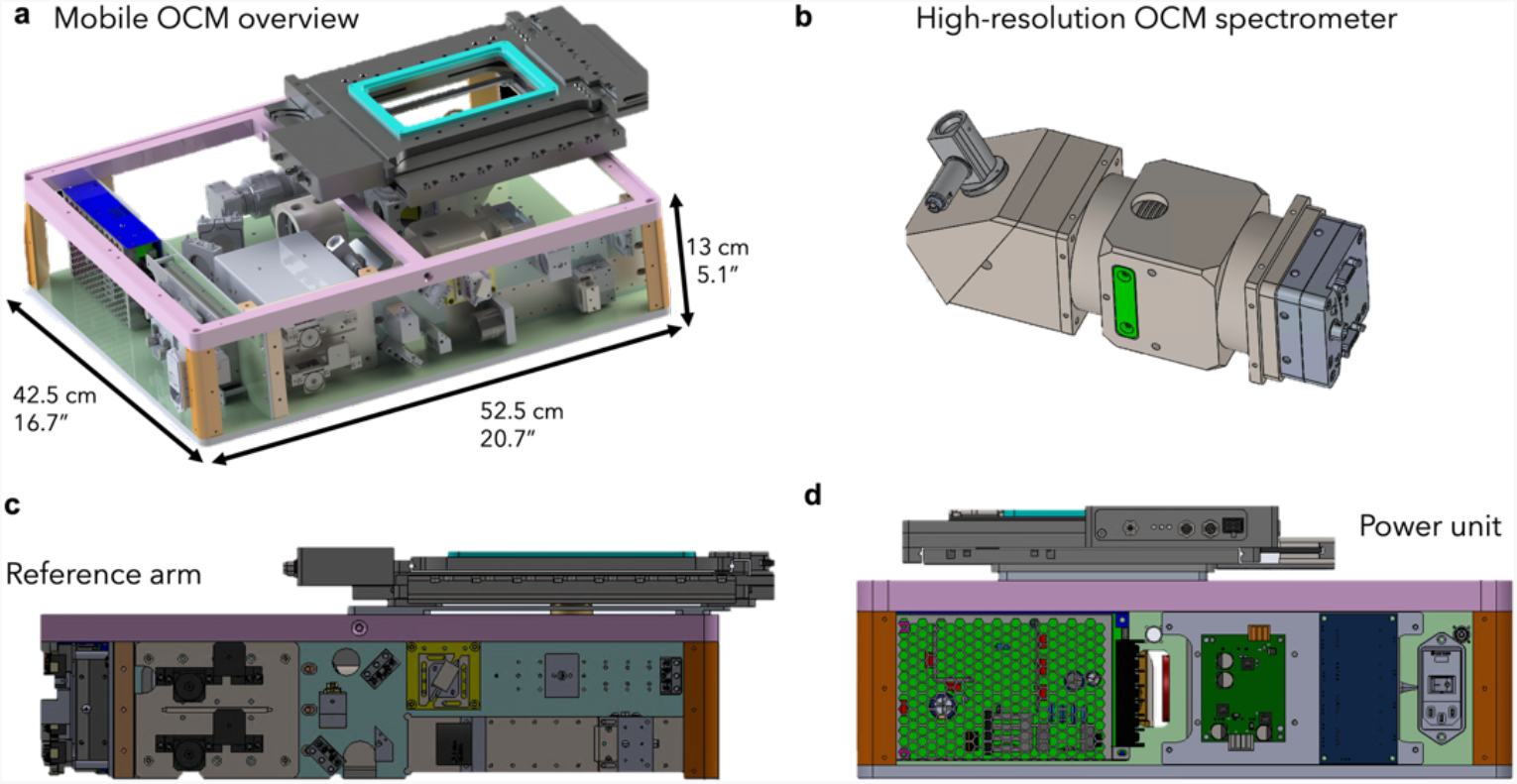
Detailed design of the mobile OCM system. **(a)** CAD rendering of the fully assembled optical module, showing the complete integration of interferometer components, galvo scanning unit, detection arm, and sample stage within the compact enclosure (42.5 × 52.5 × 13 cm). The cyan-highlighted region indicates the motorized XY sample scanning stage; the blue component (left) houses the electronics and FPGA acquisition board. **(b)** Close-up CAD view of the custom high-speed spectrometer housing with the line camera. **(c)** Side view of the mobile OCM, showing the interferometer components including the fiber coupler and reference arm optics, as well as the dispersion compensation prism pair. **(d)** Front view showing the electronics and power supply, illustrating the fully self-contained architecture of the mobile platform. Also see **Supplementary Video 5**.

## Notes

### Competing Interest Statement

The authors have declared no competing interest.

